# *HiCAGE* : an R package for large-scale annotation and visualization of 3C-based genomic data

**DOI:** 10.1101/315234

**Authors:** Michael J. Workman, Tiago C. Silva, Simon G. Coetzee, Dennis J. Hazelett

## Abstract

Chromatin interactions measured by the 3C-based family of next generation technologies are becoming increasingly important for measuring the physical basis for regulatory interactions between different classes of functional domains in the genome. Software is needed to streamline analyses of these data and integrate them with custom genome annotations, RNA-seq, and gene ontologies. We introduce a new R package compatible with Bioconductor—Hi-C Annotation and Graphics Ensemble (*HiCAGE*)—to perform these tasks with minimum effort. In addition, the package contains a shiny/R web app interface to provide ready access to its functions.

**Availability and Implementation:** The software is implemented in R and is freely available under GPLv3. *HiCAGE* runs in R (version ≥3.4) and is freely available through github (https://github.com/mworkman13/HiCAGE) or on the web (https://junkdnalab.shinyapps.io/hicage).

## Introduction

The three-dimensional organization of the genome plays a fundamental role in the regulation of gene expression [4]. In the last 15 years, Chromosome Conformation Capture (3C) technology [5] has enabled a detailed exploration of nuclear architecture by identifying regions of interacting DNA. Subsequent 3C-based methods (*e.g.* 4C [14], 5C [6], Hi-C [11], ChIA-PET [8], and HiChIP [12] have allowed for more comprehensive and precise analysis of chromatin interactions by incorporating next-generation sequencing and chromatin immunoprecipitation into the original method. Multiple browsers are available for exploration of 3D-interactions in the context of genomic annotations [11, 15]. It is currently possible to annotate enhancer-promoter interactions with R [10]). However, comprehensive annotation of interacting genomic regions generated from 3C-based data integrated with gene expression analysis and gene ontology is lacking. Here we introduce an R package *HiCAGE* which provides these functions, as well as graphical summaries of integrations and gene-ontology enrichment of candidate genes based on proximity.

## Software Overview

The *HiCAGE* (Hi-C Annotation and Graphics Ensemble) package annotates and visualizes interacting chromatin regions. *HiCAGE* integrates processed interaction regions derived from 3C-based data with genomic annotations and assigns genes by proximity. Gene expression data are optionally integrated. *HiCAGE* filters expression lists by interaction class (*e.g.* “promoter-enhancer”). It also enables subsequent gene-ontology analysis of these filtered lists. Importantly, *HiCAGE* outputs a data frame with unique annotations for each interaction region and its nearest gene, a challenging and critical step for downstream analysis.

### Data Import

The *HiCAGE* package imports data from tab-delimited text files. 3C-based data includes chromosome, genome coordinates, and interaction score or q-value of the loop. The package includes a sample dataset of processed Hi-C [13] containing loop annotations of K562 cells, a *StatePaintR* [2] track, and an RNA-seq gene expression file from the ENCODE project [7]. Chromatin state annotations are currently available through *StateHub*/*StatePaintR* (www.statehub.org) for the ENCODE cell lines, RoadMap tissues, CEEHRC cells and tissues, and Blueprint blood cells. Any generic chromatin state annotations similarly formatted may be used. Data is input and processed in *HiCAGE* with or without gene expression data. Additionally, data-containing columns are user-defined for each of the data sets to allow flexibility of input files.

### Integration of Expression Profiles

Each region in the 3C-based data is annotated with overlapping segmentation marks contained in the annotation (bed format, see example data) file. *HiCAGE* utilizes the ranking of *StatePaintR* [2] to prioritize states for each genomic segment and assigns a state for each region. Once chromatin states have been defined, the expression values from the RNA-seq file are associated with the nearest gene.

*HiCAGE* generates an object containing the chromosome and coordinates of the interacting regions, a chromatin state and rank score for each region, nearest gene and expression if an RNA-seq file was input, and the interaction statistic. Data are summarized in a table, which can be filtered for genes associated with each defined segmentation mark that interacts with a given distal mark (*e.g.* all genes associated with active promoters interacting with active enhancers).

### Gene Ontology Analysis

*HiCAGE* streamlines analysis of Gene Ontology enrichment with the *TopGO* package [1]. The user selects a category of interacting annotations (interaction pairs) for enrichment analysis, and associated genes are evaluated relative to the background of genes associated with all other interaction pairs. Users may filter genes to remove low- or non-expressed genes. Enrichment statistics are produced for each significant term. By default, terms are ordered by classic Fisher’s exact-test *p*-value.

### Summary and Visualization

To capture the complex relationship between genomic architecture, chromatin state interaction, and gene expression on a whole genome level, *HiCAGE* utilizes the R package *Circlize* [9] to integrate the three datasets visually as shown in Figure 1a. This representation makes it possible to assess at a glance which chromatin states interact most frequently, which have the highest interaction scores, and which are associated with the highest gene expression. Each plot diagrams the number of interactions between each chromatin state. Average interaction score between each interaction pair is shown on a concentric heatmap surrounding the circos plot. If gene expression data was input, gene expression is displayed on the outermost ring. *HiCAGE* provides visual summaries of links between different annotations in *UpSetR* [3] plots (Figure 1b). Using the genelist tab in *HiCAGE* we selected the interaction pair ‘enhancer-promoter’, and performed gene ontology analysis. The most enriched terms were ‘lamin filament’ (*p* = 0.0020, Fisher’s exact test) and ‘blood microparticle’ (*p* = 0.0046).

**Figure 1.**
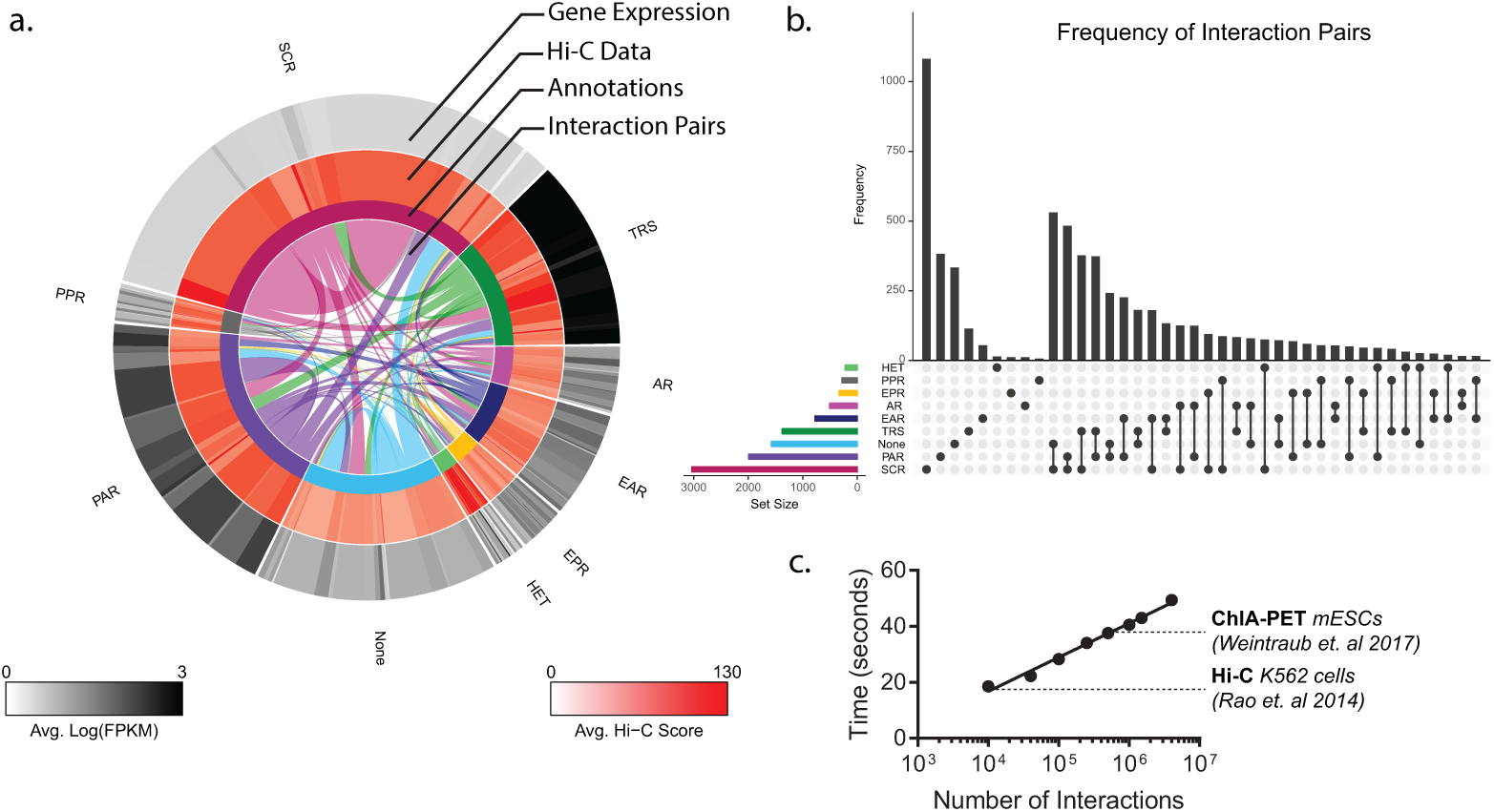
a. Example using Hi-C data from the K562 cell line [13]. The inner set of links illustrates by link width the number of connections between each pair of categories. Each type of pair is represented twice to cover both possible orientations along the chromosome. Abbreviations (clockwise from top): SCR—Epigenetically silenced chromatin; TRS—transcriptionally active regions; AR—Active region, EAR— Enhancer, active; EPR—Enhancer, poised; HET—Heterochromatin; None—no histone marks/annotations associated; PAR—Promoters, active; PPR—Promoters, poised. b. Summary of link frequencies. UpSetR view of the frequency of links between each pair of annotation classes. c. Computation time as a function of number of interactions.

Both the circular and annotation summaries can be generated dynamically on the shiny app. To illustrate these concepts and validate *HiCAGE*, we plotted data for YY1 and CTCF ChIA-pet from [16]. This study demonstrated that in contrast to the DNA-binding insulator protein CTCF, which demarcates the boundaries of topological association domains, YY1 mediates local interactions within these domains. *HiCAGE* analysis of these data confirms these results, showing that for YY1, Promoter-promoter and promoter-enhancer interactions predominate (Supplementary Figure 1a, b) whereas for CTCF, homotypic interactions between CTCF and CTCF and None annotations predominate (Supplementary Figure 1c, d).

## Conclusions

As we begin to develop a better understanding of the relationship between chromatin architecture and gene expression, more advanced tools are needed to process and visualize the data. The *HiCAGE* package is the first, to our knowledge, whole genome annotation and visualization tool for integration of NGS data types with 3C technology-based data. The plots generated by the package can be used to visualize reorganization of chromatin structure and gene expression changes on a whole genome level in response to a given stimulus or differentiation. Additionally, the package presented here will increase our understanding of the functional consequences of changes to the nuclear architecture by linking gene expression with chromatin state interactions.

## Data Sources

K562 Hi-C hg19 (GEO: GSE63525)

K562 RNA-seq [7] hg19 (ENCODE: ENCFF515MUX)

K562 annotation model hg19 (StateHub: 581ff9f246e0fb06b4b6b178 “default”)

mESC YY1 & CTCF ChIA-PET liftover to mm10 (GEO: GSE99521)

mESC RNA-seq mm10 (ENCODE: ENCFF243VGH)

es-bruce4 unknown mm10 annotation model (StateHub: 581ff9f246e0fb06b4b6b178 “default”)

## Acknowledgements

The study was funded by National Institutes of Health UO1CA184826 awarded to Benjamin Berman and RO1CA190182 (DJH).

